# Intracerebral transfection of anti-rabies virus antibodies is an effective therapy for rabies

**DOI:** 10.1101/667949

**Authors:** Washington C. Agostinho, Paulo E. Brandão

## Abstract

Rabies is a zoonotic neurological disease with 100% lethality. Some of the rare human patients who survived after multiple drug treatment have inherited severe sequelae. The objective of this study was to investigate the action of the transfection of antibodies against rabies in the central nervous system of mice as target therapy for rabies.

**Author summary:** The present study showed that after 48 h of RABV inoculation, mice injected by the intracerebral route with anti-RABV F(ab’)_2_ complexed with Bioporter® Protein Delivery Reagent (Genlantis) as a transfection agent, showing a morbidity/mortality rate of 30% with a minimum incubation period of seven days, while in the control group a significantly higher (p<0.0198) 90% morbidity/mortality was reached in thirteen days after a maximum 5-day incubation period, suggesting that the transfection of anti-RABV antibodies into the brain might prevent or delay RABV dissemination in an early stage of rabies infection. For the first time, a single compound was able to inhibit replication of the virus in the nervous system with high efficiency. This result can provide important results for the planning of protocols to prevent the fatal outcome of the disease in advanced stages. New studies focusing on the optimization of intracellular antibody delivery may be one of the main bases for more effective anti-rabies therapy.

## Introduction

Rabies is a zoonotic neurological disease with 100% lethality; some of the few human patients who survived after a multi-drug treatment developed severe motor impairment due to ischaemic encephalopathy followed by necrosis of the hippocampus, cerebellum and cortex [1–2]. The use of immunomodulators and antivirals has not been shown to be effective in inhibiting the progression of the disease when tested in mice and humans [3–4]. Circa 59,000 human deaths occur worldwide yearly due to rabies, and although it is preventable with pre and post-exposure prophylaxis, the logistics and costs involved in rabies treatments are a limiting factor to saving lives [5].

Rabies lyssavirus RABV (*Mononegavirales*: *Rhabdoviridae*: *Lyssavirus*) is a neurotropic virus with a circa 11 Kb negative-sense single-stranded RNA as a genome that codes for the nucleoprotein (N), phosphoprotein (P), matrix protein (M), envelope glycoprotein (G) and the RNA-dependent RNA-polymerase L protein [6], and it is most often transmitted amongst mammals via saliva after an initial local replication in muscle cells that follows to the central nervous system (CNS) via axons [7]. Within a variable period of time after infection, signs of hyperactivity, hypersalivation and hydrophobia are detectable. The virus causes enough damage to the brain in a few days that the infection invariably leads to coma and death by cardio-respiratory arrest [8].

Here, we show that the use of intracerebral transfection of anti-RABV antibodies to treat mice inoculated with RABV reduces mortality and extends the incubation period of rabies.

## Results

Probing the transfection to mouse brains with Bioporter agent complexed with an FITC-antibody control protein (Genlantis) after intracerebral inoculations in mice showed fluorescent foci at 4 and 6-hours post-injection in brain slices obtained in a cryomicrotome (Fig 1), evidencing the efficacy of the protein transfection to mouse brains. However, the fluorescence technique performed with microscopic slide tissue fragments showed absence of fluorescent foci using equine anti-IgG conjugate for the mice inoculated with the F(ab’)_2_ anti-RABV Bioporter complex at concentrations of 50 and 250 μg/mL and Bioporter, for post-inoculation periods of 4 and 6 h (Unpublished data).

**Fig. 1.**
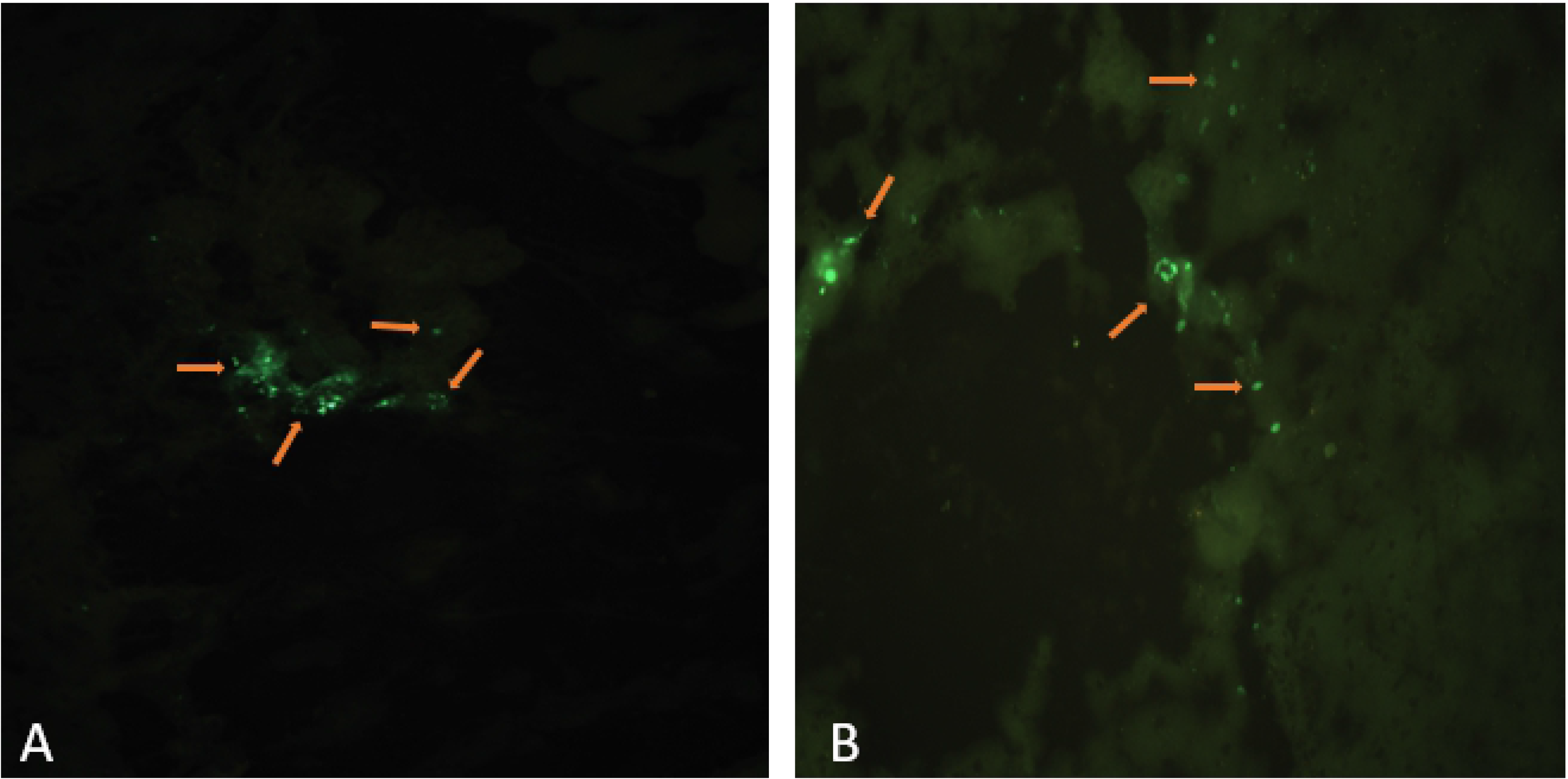
Ten-μm cryosections of mice brain after transfection of antibodies with FITC-antibody control protein® (Genlantis) as a transfection tracer in the central nervous system of mice using Bioporter® (Genlantis) as a transfection agent showing fluorescent dots (arrows) on the cytoplasm at 4 (A) and 6 (B) hours post-injection. 200× increase.

Mice which received anti-RABV F(ab’)_2_ in conjunction with Hepes 48 h,p.i., the mice in the control group had onset of rabies symptoms at 5 days post injection. About 90% of the mice showed symptoms of 2 to 3 days, followed by death, with only one mouse showing no symptoms throughout the experiment, resulting in 90% mortality. Only one mouse presented symptoms late, at 9 days p.i., consolidating survival of 10%. The clinical signs observed were anorexia, piloerection, arching of the back, and limb paralysis before death.

On the other hand, the morbidity/mortality rate in the group treated with Bioporter plus anti-RABV F(ab’)_2_ was as low as 30% with a minimum incubation period of seven days, resulting in a significant difference (p<0.0198) when compared to the control group (Fig 2).

**Fig. 2.**
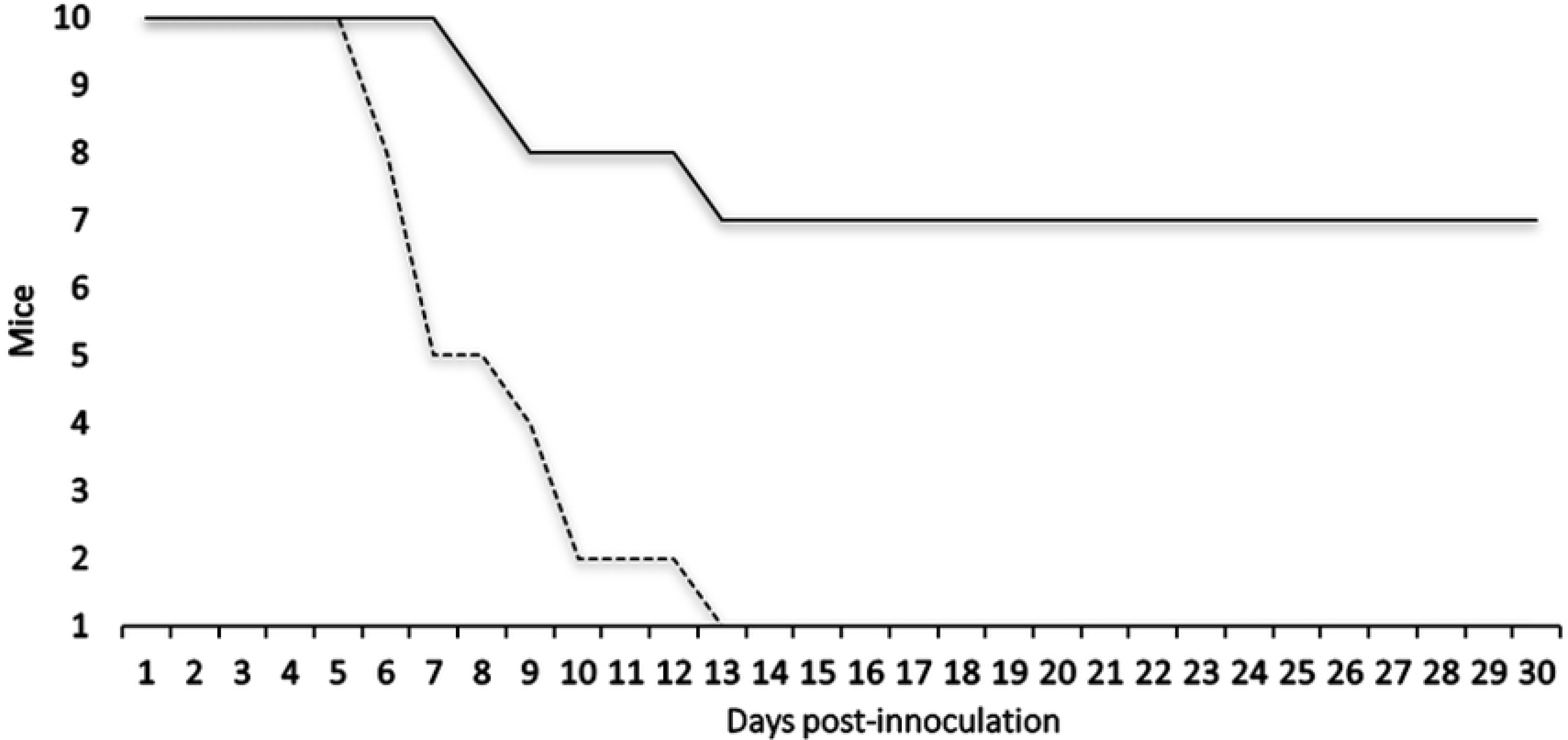
Survival plot for mice inoculated intracerebrally with 10^3.8^ DL_50%_RABV DOG-IP3629/11 and treated 48 h post-inoculation with Anti-RABV F(ab’)_2_ plus Bioporter ® (solid line) or Anti-RABV F(ab’)_2_ plus Hepes 20 mM pH 7.4 (dashed line).

Bioporter alone had no significant action on RABV, as morbidity/mortality rates of 50 and 80% were found for mice treated with only Bioporter or Hepes 20 mM pH 7.4 solution, respectively, after 48 h of RABV inoculation (p=0.3498), indicating that the reduced morbidity/mortality rate in mice treated with anti-RABV F(ab’)_2_ transfected with Bioporter was due to a specific intracellular neuralization effect (Unpublished data).

All dead animals were positive for direct immunofluorescence for RABV, with no difference in fluorescence intensity between groups. Thirty days after viral inoculation, all surviving mice were euthanized and negative by direct immunofluorescence and PCR for RABV.

In summary, these results show that the transfection of anti-RABV antibodies into the brain might prevent or delay RABV dissemination in an early stage of rabies infection.

## Discussion

In this study, the IFD technique used with equine anti-IgG conjugate to detect a transfection of anti-RABV F(ab’)_2_ through the Bioporter reagent performed 4-6 h after its intracerebral injection showed no fluorescence, which may indicate that (a) transfection of anti-RABV F(ab’)_2_ occurred with low efficiency, (b) its intracytoplasmic dispersion avoids large clusters of anti-RABV F(ab’)_2_ accumulators, in this way the fluorescence of the conjugate anti-IgG antibody is inhibited by fluorescence microscopy if it is associated with anti-RABV F(ab’)_2_ and (c), the result may be related to lack of affinity of the equine anti-IgG conjugate by the fragmentation of IgG- RABV.

If the latter is the case, the absence of fluorescence should result from the purification of anti-RABV F(ab’)_2_ by the enzymatic digestion of pepsin, which produces two F(ab’)_2_ fragments bound by disulfide bond, which reduces its molecular weight from 160 kDa IgG to 90 to 100 kDa, eliminating from the molecule the Fc fraction responsible for complement activation by the classical route.

Interestingly, in the work done by Weiil, among the three antibodies transfected in vitro by the Pulsin reagent, the only one that did not demonstrate the expected signaling was the Anti-mouse IgG antibody, since a secondary antibody rapidly exuded from the cytoplasm when the cells were treated with digitonin (lipid solubilizer) revealing that it was not bound to any target, while the primary antibodies remained within the cytoplasm for 15 minutes. This result helps to support hypothesis (b) in which the dispersion of anti-RABV F(ab’)_2_ in the cytoplasm, implies non-visualization of its location by signaling by the equine anti-IgG conjugate if it is associated [9].

A fluorescence microscopy depends on detection of the fluorophore above its effective detection limit. And, eventually, false negatives may occur when attempting to study the dispersion of fluorophore labeled molecules, so that a low level of fluorescence could reflect the absence in that tissue area or a high degree of dispersion would decrease the fluorescence to the point of not being distinguished of the autofluorescence of the tissue attached to the slide[10]. However, the abundance of FITC-control Bioporter, the product of transfection, enabled confirmation of delivery by the reagent, even if no specific target was found intracellularly.

Since the efficacy of post-exposure treatment decreases progressively when initiated late, antibody performance is still significant if treatment is performed in the first 24 h. At this early stage, treatment with antibodies performed directly in the CNS may still prevent or delay the spread to the rest of the brain. However, in later stages, no protection is effective because the virus has spread to larger parts of the brain [11].

However, inversely, our treatment performed 48 hours later with the Bioporter reagent, 70% of the mice in the treated group were healthy after viral infection, suggesting intense inhibition of the virus by anti-RABV F(ab’)_2_ and, in addition, there was an increase in survival of 48 h for the rest of the mice that died. The anti-RABV F(ab’)_2_ was transfected using a single intracerebral inoculation with optimal result. These results are consistent with previous observations showing inhibition of viral activity in N2A cell culture inoculated with isolates DOG-IP3629/11 [12]. Therefore, transfection of anti-RABV F(ab’)_2_ *in vivo*, demonstrated in this study could act in other variants with the same result confirmed for (IP3629/11 dog), especially in cases of human rabies.

The possibility of the Bioporter reagent contributing to the antiviral action together with the anti-RABV F(ab’)_2_ required further investigation. When this possibility was tested, the Bioporter reagent was inoculated into mice in the CNS without addition of anti-RABV F(ab’)_2_. This approach led to 50% and 20% survival between the Bioporter and Hepes salts respectively, without significant difference (p=0,3498), demonstrating that the inhibitory effect of treatment was due only neutralizing action of anti-RABV F(ab’)_2_.

Infections in the CNS are contained by the action of several immune effectors such as antibodies, cytotoxic T-cells and soluble factors that are involved in generation and control of the immune response as type-1 IFNs. Consequently, after brain infection by a pathogen, MHC II expression is surprisingly upregulated by approximately 90% by glial cells, including in diffuse areas distal to viral infection [13]. In murine brain infection demonstrates the prolonged activation of microglia associated with the continued presence of long-lasting memory T cells in the brain. Thus, it is clear that this small number of long-term memory T-cells may advance control of reinfection or reactivation of pathogens in the CNS by directing innate immune cells as microglia [14].

In some reports, the serological status of the patients shows that the production of antibodies plays a fundamental role in viral clearance. Prior to the Milwaukee Protocol, few human cases of rabies survival received post-exposure prophylaxis only with administration of the vaccine [15, 16, 17, 18]. Complete recovery or with sequels in rabies patients is limited to a few cases in the literature linked to the history of immunization with the combination of vaccines and passive immunization of antibodies at the onset of symptoms [19, 20, 21]. Studies with B cell-deficient mice, which underwent CNS virus inhibition after peripheral administration of RABV-specific antibodies at 5 d.p.i, support these results. Because, passively administered antibodies gain access through the BBB, conferring therapeutic antiviral effects on the CNS. This indicates that neutralizing antibodies may be able to cross the blood-brain barrier, representing an increase in the patient’s expected life expectancy until immunotherapy can establish the response against the virus [22, 23].

Anti-RABV antibody transfection as shown in this report is a candidate new tool for the treatment of rabies when the disease has already manifested, such as in cases in which no post-exposure prophylaxis was applied or in cases the prophylaxis failed. We have added a new tool for manipulation in the against rabies, by inhibiting replication and viral synthesis in neural cells without affecting neurotransmitters, using a single intracerebral inoculation with optimal results.

## Materials and Methods

### Ethics

This experiment was approved by the Ethics Committee on Animal Use (CEUA) of the School of Veterinary Medicine and Animal Science - University of São Paulo (FMVZ - USP), under Protocol no. 9658071016. All mice were used for the experiments; prior to any procedure, mice were anesthetized with isofluoran.

### Transfection test with Bioporter® protein delivery reagent *in vivo*

First, in order to assess the effectivity of protein delivery to mice brains, a total of 40 μl of FITC-antibody control protein (Gelantis) was complexed with the Bioporter® Protein Delivery Reagent (Genlantis) per the manufacturer’s instructions and was injected by the intracerebral route in two mice, and the CNS of each mouse was collected at 4 and 6 hours post-injection. Ten-μm sections were obtained in a LEICA CM 1860 UV microtome, fixed on glass slides with −20°C acetone for 2 hours and observed for fluorescence with an OLYMPUS ® BX53 epifluorescence microscope.

### Transfection test with anti-RABV F(ab’)_2_ *in vivo*

Next, forty-two female 21-day-old CH3 ROCKEFELLER mice were inoculated with RABV strain IP3629/11-AgV2 isolated from a dog in Brazil on mouse central nervous system (CNS) with a titer of 10^3.8^ DL_50_/ μL kindly provided by the Pasteur Institute, Brazil. After 48 h, mice were intracerebrally injected with 40 μL of either Bioporter resuspended in Hepes 20 mM pH 7.4 containing anti-RABV F(ab’)_2_ (treated group, n=10 mice) or Hepes 20 mM pH 7.4 containing anti-RABV F(ab’)_2_ (control group, n=10 mice), both groups with 0.17 UI (250 μg) of anti-RABV F(ab’)_2_ as a final dose.

### Evaluation of the action of the transfection agent alone on RABV

To assess whether the Bioporter transfection agent alone had any effects on RABV, the aforementioned experiment was repeated, but the (test group, n=10 mice) was injected with 40 μL of Bioporter in 400 μl Hepes 20 mM pH 7.4, and the (control group, n=10 mice), was injected with 40 μL of Hepes 20 mM pH 7.4 solution after 48 h of RABV inoculation.

### Antibodies

Anti-rabies hyperimmune serum was kindly provided by FUNED (Fundação Ezequiel Dias), Brazil, containing enzimatic-digestion purified equine IgG Fab fragment against the PV strain of RABV (200UI/mL).

### Equine anti-igG conjugate

For evaluation of transfection of equine F(ab’)_2_ against RABV by direct immunofluorescence, Anti-Horse IgG (whole molecule) -FITC antibody (Sigma-Aldrich) was used.

### Virus

Strain (IP3629/11-AgV2 dog isolated from Brazil), grown on mice central nervous system (CNS) with a title of 10^3.8^ DL_50_/ μL, kindly provided by the Pasteur Institute, Brazil, was used for the infection in mice.

### Antibody transfection test

FITC-antibody control protein (Gelantis) complexed with Bioporter® Protein Delivery Reagent (Genlantis) per manufacturer’s instructions were injected by the intracerebral route in two mice and the CNS of each mouse was collected at 4 and 6 hours post-infections; 10μm sections were obtained in a LEICA CM 1860 UV microtoime, fixed in glas sliced with −20°C acetone/2 hours and observed for fluorescence with a OLYMPUS ® BX53 epifluorescence microscope.

### Direct fluorescent antiboy test (DFAT) and PCR

All mice were observed for period 30 days after RABV inoculation for signs of rabies such as anorexia, piloerection hyperesthesia, aggressiveness, paralysis and death, being the surviving mice euthanized at the end of the observation period. The central nervous system (CNS) of each mouse was tested with a direct fluorescente antiboy test (DFAT) [24], using an anti-RABV nucleocapsid IgG conjugates with Fluorescein isothiocyanate (Pasteur Institute, Brazil), and, if negative, to a PCR targeting RABV N-P genes [25].

### Statistical analysis

GraphPad Prism was used for statistical analyses of *in vivo* data. Fisher’s test with *α* = *0.05* was used for the statistical analysis with Fisher Exact Test Calculator online.

## Acknowledgments

The authors are thankful for the financial support provided by CNPq (Brazilian National Board for Scientific and Technological Development) grant number 307291/2017-0 and CAPES (Coordenação de Aperfeiçoamento de Pessoal de Nível Superior, Brazil) Finance Code 001, which had no role in the study design, collection, analysis and interpretation of data, writing of the report and in the decision to submit the article for publication.

## References

1. Burton EC, Burns DK, Opatowsky MJ, El-Feky HW, Fischbach B, et al. Rabies encephalomyelitis: Clinical, neuroradiological, and pathological findings in 4 transplant recipients. Arc. Neurol. 2005;62(6) 873–882. https://doi.org/10.1001/archneur.62.6.873.

2. Manoj S, Mukherjee A, Johri S, Kumar K.V. Recovery from rabies, a universally fatal disease. Mil. Med. Res. 2016; 3(21). https://doi.org/10.1186/s40779-016-0089-y

3. van Thiel PP, de Bie RM, Eftimov F, Tepaske R, Zaaijer HL, et al. Fatal human rabies due to Duvenhage virus from a bat in Kenya: failure of treatment with coma-induction, ketamine, and antiviral drugs. PLoS Negl. Trop. Dis. 2009; 3(7): e428. https://doi.org/10.1371/journal.pntd.0000428

4. Marosi A, Dufkova L, Forró B, Orsolya F, Erdélyi K, et al. Combination therapy of rabies-infected mice with inhibitors of pro-inflammatory host response, antiviral compounds and human rabies immunoglobulin. Vaccine. 2018. https://doi.org/10.1016/j.vaccine.2018.05.066

5. Hampson K, Coudeville L, Lembo T, Magangaet S, Alexia Kieffer, et al. Estimating the global burden of endemic canine rabies. PLoS Negl Trop Dis. 2015; 9(4): e0003709. https://doi.org/10.1371/journal.pntd.0003709

6. Jackson, A.C., Wunner W.H. Rabies, second ed. Academic Press, London, 2007.

7. Lewis P, Fu Y, Lentz TL. Rabies virus entry at the neuromuscular junction in nerve-muscle cocultures. Muscle Nerve. 2000; 23(95): 720–730.

8. WHO. Expert consultation on rabies: second Report. Geneva: World Health Organization, 2013.

9. Weill CO, Biri S, Erbacher P. Cationic lipid-mediated intracellular delivery of antibodies into live cells. Biotechniques. 2008;44(7):7–11.

10. Ho JK, White PJ, Pouton CW. Self-Crosslinking Lipopeptide/DNA/PEGylated Particles: A New Platform for DNA Vaccination Designed for Assembly in Aqueous Solution. Mol Ther Nucleic Acids. 2018;12(September):891–2. Available from: https://doi.org/10.1016/j.omtn.2018.05.025.

11. Terryn S. Development and evaluation of antiviral immunoglobulin single variable domains for prophylaxis of rabies in mice. PLoS Negl Trop Dis. 2016. Available from: https://doi.org/10.1371/journal.pntd.0004902

12. Kawai, JGC. Transfection of anti-rabies antibodies into n2a lineage cells (mouse neuroblastoma) and intracellular neutralization for use as an antiviral. Postdoctoral Research Report. Faculty of Veterinary Medicine and Animal Science, University of São Paulo. 2012. https://bv.fapesp.br/en/auxilios/31673/transfeccao-de-anticorpos-anti-ravas-virus-in-cellular-cells-n2-neuroblastoma-of-camundong/

13. Hooper DC, Phares TW, Fabis MJ, Roy A. The production of antibody by invading B cells is required for the clearance of rabies virus from the central nervous system. PLoS Negl Trop Dis. 2009;3(10):1–8.

14. Prasad S, Lokensgard JR. Brain-Resident T Cells Following Viral Infection. 2019;32(1):48–54.

15. Hattwick MAW, Weis TT, Stechschulte CJ, Baer GM, Gregg MB. Recovery from Rabies. N Engl J Med [Internet]. 1972;(76):931–42. Available from: http://www.nejm.org/doi/abs/10.1056/NEJMe058092

16. Porras C, Barboza JJ, Fuenzalida E, Adaros HL, Bioch Amo de D, Furst J. Recovery From Rabies in Man. JAMA J Am Med Assoc. 1966;(85):44–8.

17. Alvarez, H. L; Fajardo; V. R.; Pedroza, R. R.; Hemachudha, T.; Kamolvarin, N.; Cortes, C. G.; Baer GM. Partial recovery from rabies in a nine-year-old boy. Pediatr Infect Dis J. 1994;13:1154–5.

18. Madhusudana SN, Nagaraj D, Uday M, Ratnavalli E, Verendra Kumar M. Partial recovery from rabies in a six-year-old girl [2]. Int J Infect Dis. 2002;6(1):85–6.

19. Karahocagil MK, Akdeniz H, Aylan O, Sünnetçioğlu M, ün H, Yapici K, et al. Complete Recovery from Clinical Rabies: Case Report. Turkiye Klin J Med Sci [Internet]. 2013;33(2):547–52. Available from: http://www.tipbilimleri.turkiyeklinikleri.com/abstract_64465.html

20. Willoughby RE, Tieves KS, Hoffman GM, Ghanayem NS, Amlie-Lefond CM, Schwabe MJ, et al.Survival after Treatment of Rabies with Induction of Coma. N Engl J Med [Internet]. 2005;352(24):2508–14. Available from: http://www.nejm.org/doi/abs/10.1056/NEJMoa050382

21. PROMED. Rabies, human survival, bat-Brazil: (Pernambuco). 2009; 14 Nov: 20081114.3599. Available from: https://www.promedmail.org/post/20090919.3292

22. Roy A, Hooper DC. Lethal Silver-Haired Bat Rabies Virus Infection Can Be Prevented by Opening the Blood-Brain Barrier. J Virol. 2007;81(15):7993–8. Available from: http://jvi.asm.org/cgi/doi/10.1128/JVI.00710-07

23. Hooper, D Craig, Roy Anirbal, Kean B Rhonda, Phares TW B DA. Therapeutic immune clearance of rabies virus from the CNS. Future Virol. 2011 Mar. 1; 6(3): 387–397. doi:10.2217/fvl.10.88.

24. Dean DJ, Abelseth MK, Atanasiu P. Fluorescent antibody test, in: Meslin FX, Kaplan MM, Koprowski H. (Eds.), Laboratory techniques in rabies. World Health Organization, Geneva, 1996. 88–95.

25. Orciari, LA, Niezgoda M, Hanlon CA, Shaddock JH, Sanderlin DW, et al. Rapid clearance of SAG-2 rabies virus from dogs after oral vaccination. Vaccine. 2001; 14;19(31): 4511–4518. https://doi.org/10.1016/S0264-410X(01)00186-4.

